# Chronic stress antagonizes formation of Stress Granules

**DOI:** 10.1101/2024.10.29.620814

**Authors:** Yuichiro Adachi, Allison M. Williams, Masashi Masuda, Yutaka Taketani, Paul J. Anderson, Pavel Ivanov

## Abstract

Chronic stress mediates cellular changes that can contribute to human disease. However, fluctuations in RNA metabolism caused by chronic stress have been largely neglected in the field. Stress granules (SGs) are cytoplasmic ribonucleoprotein condensates formed in response to stress-induced inhibition of mRNA translation and polysome disassembly. Despite the broad interest in SG assembly and disassembly in response to acute stress, SG assembly in response to chronic stress has not been extensively investigated. In this study, we show that cells pre-conditioned with low dose chronic (24-hour exposure) stresses such as oxidative stress, endoplasmic reticulum stress, mitochondrial stress, and starvation, fail to assemble SGs in response to acute stress. While translation is drastically decreased by acute stress in pre-conditioned cells, polysome profiling analysis reveals the partial preservation of polysomes resistant to puromycin-induced disassembly. We showed that chronic stress slows down the rate of mRNA translation at the elongation phase and triggers phosphorylation of translation elongation factor eEF2. Polysome profiling followed by RNase treatment confirmed that chronic stress induces ribosome stalling. Chronic stress-induced ribosome stalling is distinct from ribosome collisions that are known to trigger a specific stress response pathway. In summary, chronic stress triggers ribosome stalling, which blocks polysome disassembly and SG formation by subsequent acute stress.

**Significant statements:** Stress granules (SGs) are dynamic cytoplasmic biocondensates assembled in response to stress-induced inhibition of mRNA translation and polysome disassembly. SGs have been proposed to contribute to the survival of cells exposed to toxic conditions. Although the mechanisms of SG assembly and disassembly in the acute stress response are well understood, the role of SGs in modulating the response to chronic stress is unclear. Here, we show that human cells pre-conditioned with chronic stress fail to assemble SGs in response to acute stress despite inhibition of mRNA translation. Mechanistically, chronic stress induces ribosome stalling, which prevents polysome disassembly and subsequent SG formation. This finding suggests that chronically stressed or diseased human cells may have a dysfunctional SG response that could inhibit cell survival and promote disease.

## Introduction

Eukaryotic cells have evolved universal stress-response programs that allow them to survive under toxic environmental conditions (1, 2). Stress granule (SG) assembly, an early response to cellular stress, is triggered by stress-induced global translation inhibition and polysome disassembly (3, 4). Canonical SGs are membrane-less cytoplasmic RNA granules consisting of 40S ribosome-contained mRNPs, translation initiation factors, mRNAs, RNA binding proteins, and signaling molecules. SG composition varies in a stress-specific manner and determines whether cells live or die (5, 6). For instance, while sodium arsenite (SA), commonly used to mimic oxidative stress, induces the assembly of canonical SGs that promote cell survival, nitric oxide induces the assembly of non-canonical SGs linked to decreased cell viability (7). SGs also contribute to the aggregation of disease-related proteins in the motoneurons of patients with neurodegenerative diseases such as amyotrophic lateral sclerosis (8). Both acute and chronic stress can contribute to disease pathogenesis (9). We and others have previously reported the effects of chronic stress on SG formation: chronic Bisphenol A exposure suppresses SG formation, chronic proteasomal inhibition impairs SG formation, and chronic glucose starvation induces pro-death SGs (10–12). However, the molecular mechanisms by which acute and chronic stress differentially modulate SG assembly and cell survival are poorly understood.

SA triggers phosphorylation of the α-subunit of eukaryotic initiation factor 2 (eIF2α) to inhibit translation initiation (6), which is a common initiator of the cellular stress response. eIF2 is a part of the ternary complex that delivers initiator tRNA (tRNA_i_^Met^) to the 40S ribosomal subunit. Together with other initiation factors, the ternary complex becomes the 43S mRNA pre-initiation complex, which scans the 5’-untranslated region of mRNA. When an optimal AUG codon is found, the 60S ribosomal subunit joins the 43S complex to form an 80S ribosome capable of translation elongation (13). Phosphorylation of eIF2α at Serine 51 prevents GDP/GTP exchange on eIF2 and ultimately inhibits rates of translation initiation. Because phosphorylated eIF2α inhibits mRNA translation initiation, this quickly causes polysome disassembly. In turn, disassembly of polysomes promotes SG formation. Like SA, many other stress inducers (e.g., ER stress or starvation stress) activate eIF2α kinases, induce translation inhibition and promote SG formation (14). Puromycin, a translation elongation inhibitor that causes premature termination of polypeptide synthesis, also inhibits translation and promotes SG formation (15). Puromycin forces 80S ribosomes to dissociate into 40S and 60S ribosomal subunits, effectively stripping ribosomes off of the mRNA. In contrast, translation elongation inhibitors such as anisomycin (ANS), cycloheximide, and emetine “freeze” ribosomes onto the mRNA, preventing untranslated transcripts from assembling SGs (15, 16). While the above drugs are known to regulate SG dynamics, how impairment of the translation elongation under physiological (unrelated to pharmacological treatment) situations affects SG formation is largely understudied.

The elongation step of mRNA translation is also well regulated. The primary target is eukaryotic elongation factor 2 (eEF2), a GTP-dependent translocase responsible for the movement of nascent peptidyl-tRNAs from the A-site to the P-site of the ribosome. Its activity is regulated by phosphorylation on threonine 56 (T56) by eEF2 kinase (eEF2K), namely causing downregulation of its activity. (17, 18). In turn, eEF2K itself is a phosphorylation target for diverse pathways to impact elongation, ranging from signaling from nutrient deprivation to changes in cell cycle. In addition, the rate of ribosome elongation can also be affected by mRNA secondary structures, an insufficient supply of aminoacyl-tRNAs, or inefficient ribosome termination/recycling, which can cause ribosomes to stall and collide (19–21).

Ribosome stalling and collisions can also be induced by oxidative stress, starvation, ultraviolet irradiation, or stress-induced mRNA damage (22–25). Collided ribosomes are translationally incompetent and quickly recognized and degraded by the ribosome-associated quality control (RQC) pathway. Extensive ribosome collision induces the ribotoxic stress response (RSR) which is initiated by phosphorylation of ZAKα kinase to activate a pro-death signaling cascade (25–28).

SGs are in dynamic equilibrium with polysomes, and this equilibrium can be affected by acute and chronic stresses. While SGs contribute to cell survival or cell death, a connection between chronic stress, SG formation and ribosome stalling under physiological conditions has not been previously well investigated. Here, we report that various chronic stressors (oxidative stress, ER stress, mitochondria stress, and starvation) render cells incompetent for SG assembly in response to acute stress. The data on polysome profiling and the phosphorylation of translation elongation factor eEF2 implicate incomplete polysome disassembly in this process. Specifically, chronic stress slows down translation elongation and promotes ribosome stalling and thus antagonizes efficient SG formation.

## Results

### Chronic stress conditioning inhibits SG formation

To understand the effects of chronic stress on SG formation, we incubated human osteosarcoma U2OS cells with various doses (10, 50, 100, 500 μM) of sodium arsenite (SA), for different times (1, 4, 12, 24 h) prior to quantifying SG assembly (**Fig. 1 A**). We also preincubated cells with low-dose of various stressors (including 10 μM SA which did not induce SG assembly) for 24 hours prior to treating with 100 μM SA (**Fig. 1 B**). In all cases, chronic stress pre-conditioning inhibited acute SA-induced SG assembly (**Sup Fig. 1** and **Fig. 1 C – F**). SA induces SG formation via phosphorylation of eIF2α, but some other SGs inducers (e.g., rocaglamide A (Roc A)) are independent of eIF2α phosphorylation. Like SA, RocA induced fewer SGs in chronic stress pre-conditioned cells (**Sup Fig. 2**). These data suggest that chronic stress pre-conditioning interferes with acute stress-induced SG assembly irrespective of the mechanism by which translation initiation is inhibited.

**Figure 1.**
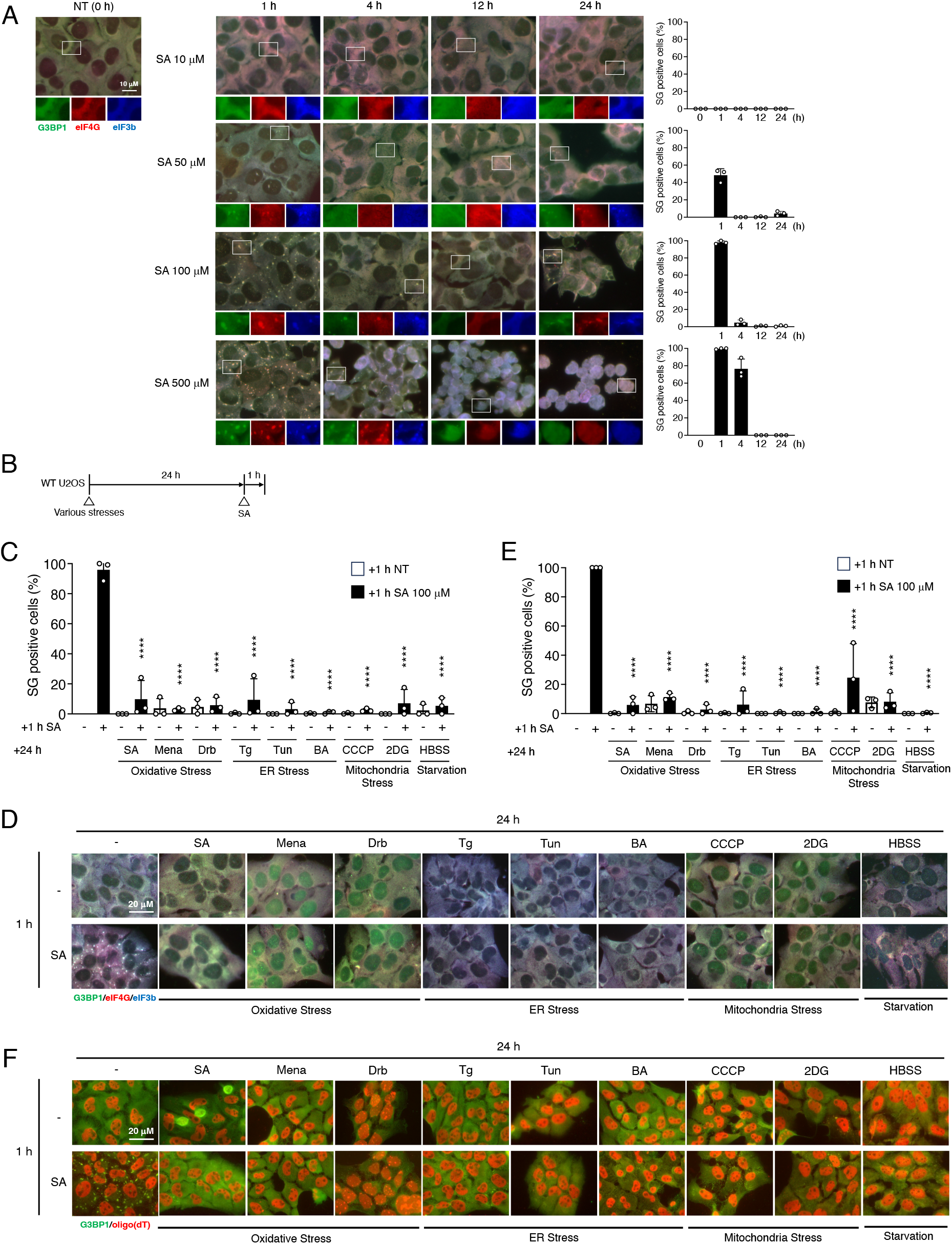
Chronic stress pre-incubated cells fail to respond to form SGs by acute stress. (A) U2OS cells were subjected to treatment with SA (10, 50, 100, 500 μM) at each time (0, 1, 4, 12, 24 h). Cells were examined for the presence of the core SG markers G3BP1 (green), eIF4G (red), and eIF3b (blue). All positive cells were quantified. Results are mean ± S.E.M. (n = 3). (B) Schematic illustration of the experimental timeline. (C–F) U2OS cells were subjected to treatment with 100 μM SA for 1 h after preincubation with 10 μM SA, 30 μM Mena (menadione), 2 μM Drb (doxorubicin), 1 μM Tg (thapsigargin), 25 μg/ml Tun (tunicamycin), 25 μg/ml BA (brefeldin A), 60 μM CCCP (carbonyl cyanide m-chlorophenyl hydrazone), 60 mM 2DG (2-deoxy-D-glucose), or HBSS for 24 h. Unstressed cells (NT) were used as a control. (C) Cells were examined for the presence of the core SG markers G3BP1, eIF4G, and eIF3B. (D) Representative images of U2OS cells stained with G3BP1 (green), eIF4G (red), and eIF3B (blue) after the cells had been subjected to specific stresses. (E) Cells were examined for the presence of the core SG marker G3BP1 and poly (A) mRNAs [FISH using oligo(dT) probe]. (F) Representative images of U2OS cells stained with G3BP1 (green) and oligo(dT) (red) after the cells had been subjected to specific stresses. (C and E) P values were assessed using a one-way ANOVA (vs. 1 h SA; *p*\*\*\*\* < 0.0001) Results are mean ± S.E.M. (n = 3).

### Incomplete polysome disassembly in chronic stress pre-conditioned cells

Since SG formation is mediated by global translation inhibition and SA induces phospho-eIF2α-dependent translational repression (29), we hypothesized that SA may not inhibit translation in chronic stress pre-conditioned cells. We compared several different types of stress (SA as an oxidative stress, Tg as an ER stress, CCCP as mitochondrial stress, and HBSS as starvation stress) in these experiments. It is evident that a 1 h incubation with 100 μM SA triggers phosphorylation of eIF2α and inhibits protein synthesis (**Fig. 2 A**). Moreover, the expression levels of SG core proteins G3BP1, G3BP2, Caprin1, and USP10 are not affected under these conditions (**Fig. 2 A**). m^7^GTP pull-downs reveal the cap-binding eIF4F complex to be intact under these conditions suggesting that mTOR signaling is not involved in the suppression of SG formation (**Sup Fig. 3**). Translation inhibition by SA rapidly disassembles polysomes (**Fig. 2 B – E**, 1h SA), which triggers efficient SG assembly (30). In contrast, 1 h incubation of 100 μM SA did not completely disassemble polysomes in chronic stress pre-conditioned cells (**Fig. 2B – E**).

**Figure 2.**
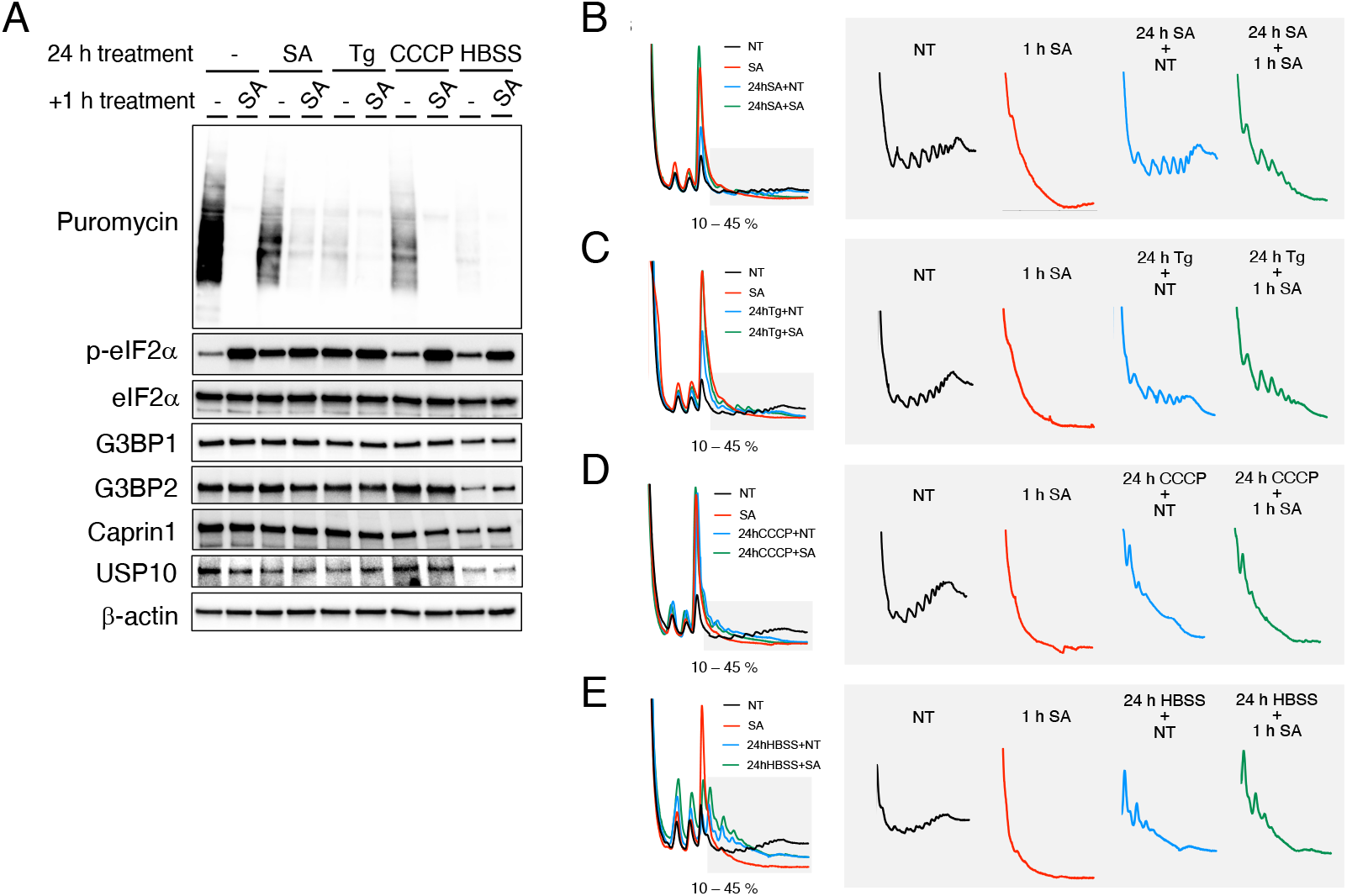
Chronic stress pre-incubated cells fail to disassemble polysomes by acute stress. U2OS cells were subjected to treatment with 100 μM SA for 1 h after pre-incubation with 10 μM SA, 1 μM Tg, 60 μM CCCP, or HBSS for 24 h. Unstressed cells (NT) were used as a control. (A) Cells were pulsed with puromycin and emetine for 5 min and lysed. Cell lysates were subjected to western blotting using antibodies for Puromycin, p-eIF2α, total eIF2α, G3BP1, G3BP2, Caprin1, USP10, and β-actin. (B–E) Polysome profiles from U2OS cells. NT; black, 1 h SA; red, (B) 24 h pre-incubation of SA; blue, 1 h SA with 24 h pre-incubation of SA; green. (C) 24 h pre-incubation of Tg; blue, 1 h SA with 24 h pre-incubation of Tg; green. (D) 24 h pre-incubation of CCCP; blue, 1 h SA with 24 h pre-incubation of CCCP; green. (E) 24 h pre-incubation of HBSS; blue, 1 h SA with 24 h pre-incubation of HBSS; green.

Because SG formation also relies on mRNA condensation besides proteins, low amounts or limited availability of mRNAs in the cells as a result of chronic stress conditions could reduce the efficiency of SG formation. Indeed, SGs are enriched in specific subsets of RNAs such as in long mRNAs (e.g., *AHNAK* and *DYNC1H1*) or long non-cording RNAs such as *NORAD* (31, 32). We thus hypothesized chronic stress decreases a pool of mRNAs and, in turn, this decrease negatively affects SGs formation. We pulled down mRNAs from total RNAs, quantified and compared the amounts of mRNAs under different conditions. mRNA pulled down efficiency was also checked by qRT-PCR, and pulled-down samples analyzed for specific mRNAs, previously shown to be enriched in SGs (**Sup Fig. 4**). The amounts of pulled-down mRNA between NT and chronic SA (10 μM) were not significantly different (**Fig. 3 A**). Additionally, the relative levels of *GAPDH, AHNAK, DYNC1H1* mRNAs and *NORAD* RNA, which are examples of efficiently enriched RNAs in SGs (31) were not significantly different (**Fig. 3 B**) thus rejecting the hypothesis that chronic stress limits pool of available RNAs for SG assembly. Moreover, a much higher dose of SA (500 μM) induces SG assembly in chronic SA (10 μM) pre-conditioned cells, consistent with the induction of polysome disassembly (**Fig. 3 C – D**). These results imply that chronic stress pre-conditioned cells retain the ability to form SGs but disassemble polysomes less efficiently, which indicates less efficient SG formation in pre-conditioned cells.

**Figure 3.**
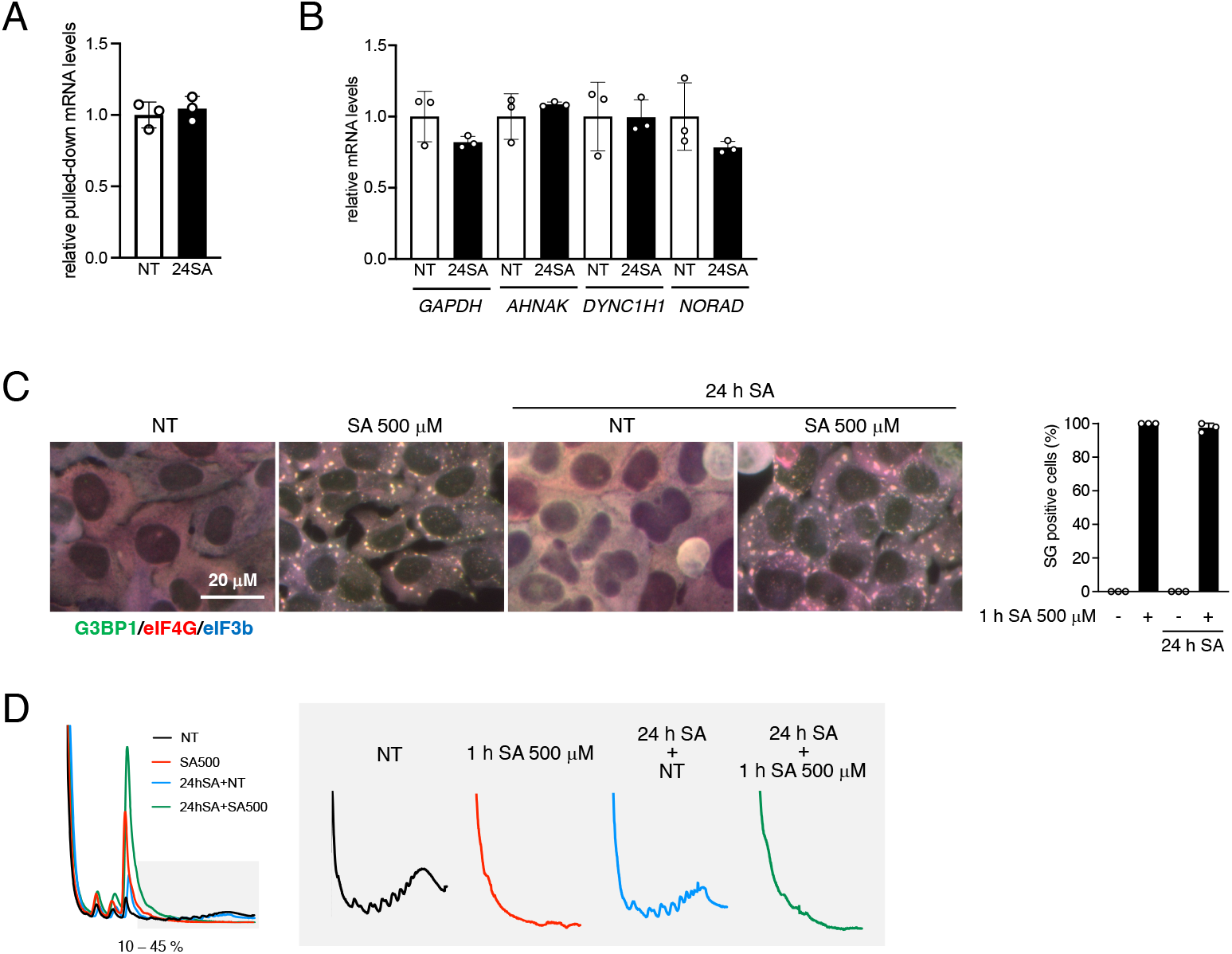
Chronic stress pre-incubated cells can form SGs by excessive stress promoting polysome disassembly. (A and B) U2OS cells were incubated with 10 μM of SA for 24 h (24SA), and total RNA was extracted. (A) mRNA was pulled down from total RNA and both RNAs were measured. Results are mean ± S.E.M. (n = 3). (B) The mRNA expression levels of GAPDH, AHNAK, DYNC1H1, NORAD were determined by qRT-PCR (scandalized by BACTIN). Results are mean ± S.E.M. (n = 3). (C and D) U2OS cells were subjected to treatment with 500 μM SA for 1 h after pre-incubation with 10 μM SA for 24 h. (C) Cells were examined for the presence of the core SG markers G3BP1 (green), eIF4G (red), and eIF3b (blue). All positive cells were quantified. Results are mean ± S.E.M. (n = 3). (D) Polysome profiles from U2OS cells. NT; black, 1 h of 500 μM SA; red, 24 h 10 μM SA pre-incubation; blue, 1 h of 500 μM SA with 24 h 10 μM SA pre-incubation; green.

### Puromycin does not promote polysome disassembly and SG formation in chronic stress pre-conditioned cells

Puromycin promotes SG assembly by inducing premature elongation termination and ribosomal subunit dissociation, while other translation elongation inhibitors such as ANS inhibit SG assembly by freezing 80S ribosomes on mRNA (**Fig. 4 A**) (15, 16, 33, 34). Consistent with these mechanisms, puromycin promotes SG assembly by an intermediate dose of SA (50 μM) in control cells (**Fig. 4B**). However, low dose (10 μM) SA pre-conditioning inhibits puromycin-induced SG assembly (**Fig. 4 B**). Polysome profiling of 50 μM SA-treated cells show the expected partial disassembly of polysomes (with ∼40% of cells being SG-positive) and complete disassembly of polysomes during puromycin co-incubation (**Fig. 4 C**). But in 10 μM SA pre-conditioned cells, puromycin does not disassemble polysomes (**Fig. 4 C** and **Sup Fig. 4**). It is known that translation elongation inhibitors such as cycloheximide freeze 80S ribosomes and inhibit SG assembly in the continued presence of stress (**Fig. 4 A**). Similar results were observed in cells treated with low dose (1 mg/ml) ANS; SA did not induce SG formation or polysome disassembly in ANS pre-incubated cells (**Fig. 4 D – E**). Our data suggest that various chronic stresses inhibit translation but do not completely disassemble polysomes (**Fig. 2**). Considering the data in both Figures 2 and 4, chronic stress may “freeze” some of the 80S ribosomes on mRNA, which inhibits ribosome dissociation and SG formation. Taken together, we hypothesized that chronic stress slows ribosome elongation.

**Figure 4.**
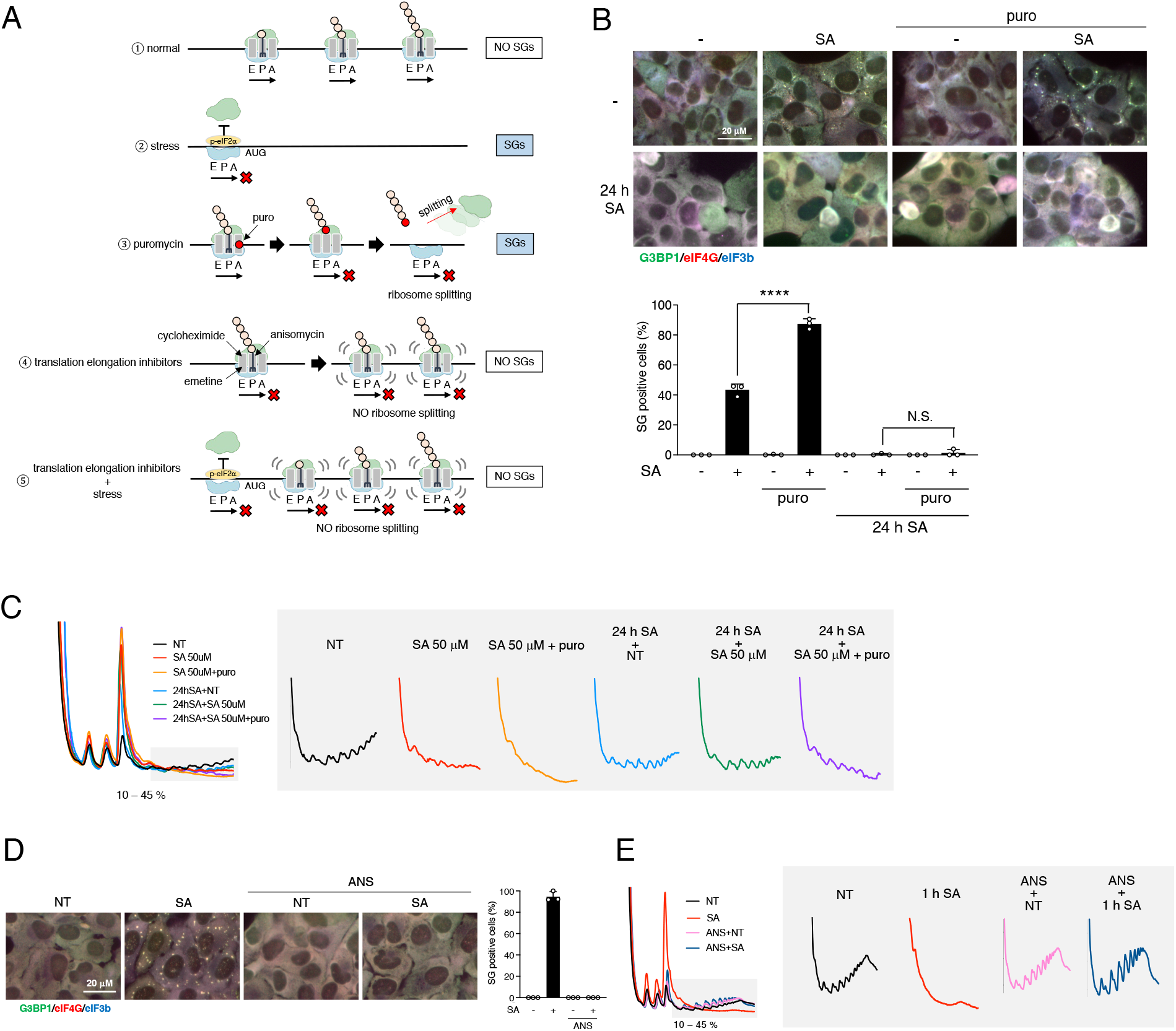
Puromycin treatment is not sufficient to promote SG formation in chronic stress pre-incubated cells. (A) Schematic illustration of the effects of puromycin and other translation inhibitors on SG formation. 1; Normal condition; 80S ribosomes are on mRNA and no SG. 2; Stressed condition; phosphorylated eIF2α (p-eIF2α) blocked to form 80S ribosomes and scanning, which prevent 80S ribosomes binding on mRNA and induces SG formation. 3; Puromycin makes ribosomes split on mRNA by synthesizing premature proteins during translation elongation, which induces SG formation. 4; Other translation elongation inhibitors freeze the translating ribosomes on mRNA, which inhibits ribosome splitting and SG formation. 5; Stress-induced eIF2α phosphorylation inhibits the initiation of translation, but translation elongation inhibitors except puromycin shown in 4 freeze translating ribosomes on mRNA and inhibit split ribosomes, which induce no SG formation. (B and C) U2OS cells were subjected to treatment with 50 μM SA for 1 h after preincubation with pre-incubation of 10 μM SA for 24 h, and 20 μg/ml puromycin (puro) for the last 0.5 h. (B) Cells were examined for the presence of the core SG markers G3BP1 (green), eIF4G (red), and eIF3b (blue). All positive cells were quantified. Results are mean ± S.E.M. (n = 3). P values were assessed using a one-way ANOVA (*p*\*\*\*\* < 0.0001, N.S.: Not Significant) (C) Polysome profiles from U2OS cells. NT; black, 1 h of 50 μM SA; red, 1 h of 50 μM SA + 0.5 h of puromycin; orange, 24 h 10 μM SA pre-incubation; blue, 1 h of 50 μM SA with 24 h SA pre-incubation; green, 1 h of 50 μM SA + 0.5 h of puromycin with 24 h SA pre-incubation; purple. (D and E) U2OS cells were subjected to treatment with 100 μM SA for 1 h after pre-incubation with 1 μg/ml ANS for 15 min. (D) Cells were examined for the presence of the core SG markers G3BP1 (green), eIF4G (red), and eIF3b (blue). All positive cells were quantified. Results are mean ± S.E.M. (n = 3). (E) Polysome profiles from U2OS cells. NT; black, 1 h SA; red, 15 min ANS; pink, 1 h SA with 15 min ANS pre-incubation; dark blue.

### Chronic stress induces translocation failure and translational delay

To determine if chronic stress slows ribosome elongation, we treated pre-conditioned cells with harringtonine, which is an inhibitor of initial elongation step of translation (35, 36). Polysome profiling showed disassembled polysomes by harringtonine incubation under normal control conditions, but harringtonine did not disassemble polysomes as efficiently (within the same time frame) in 10 μM SA chronic-stressed cells (**Fig 5. A**). This indicates that while chronically stressed cells are still translating in a manner sufficient for cell survival, the rates with which ribosomes perform mRNA translation are likely down-regulated. Such elongation slow down may also cause ribosome stalling and collisions.

**Figure 5.**
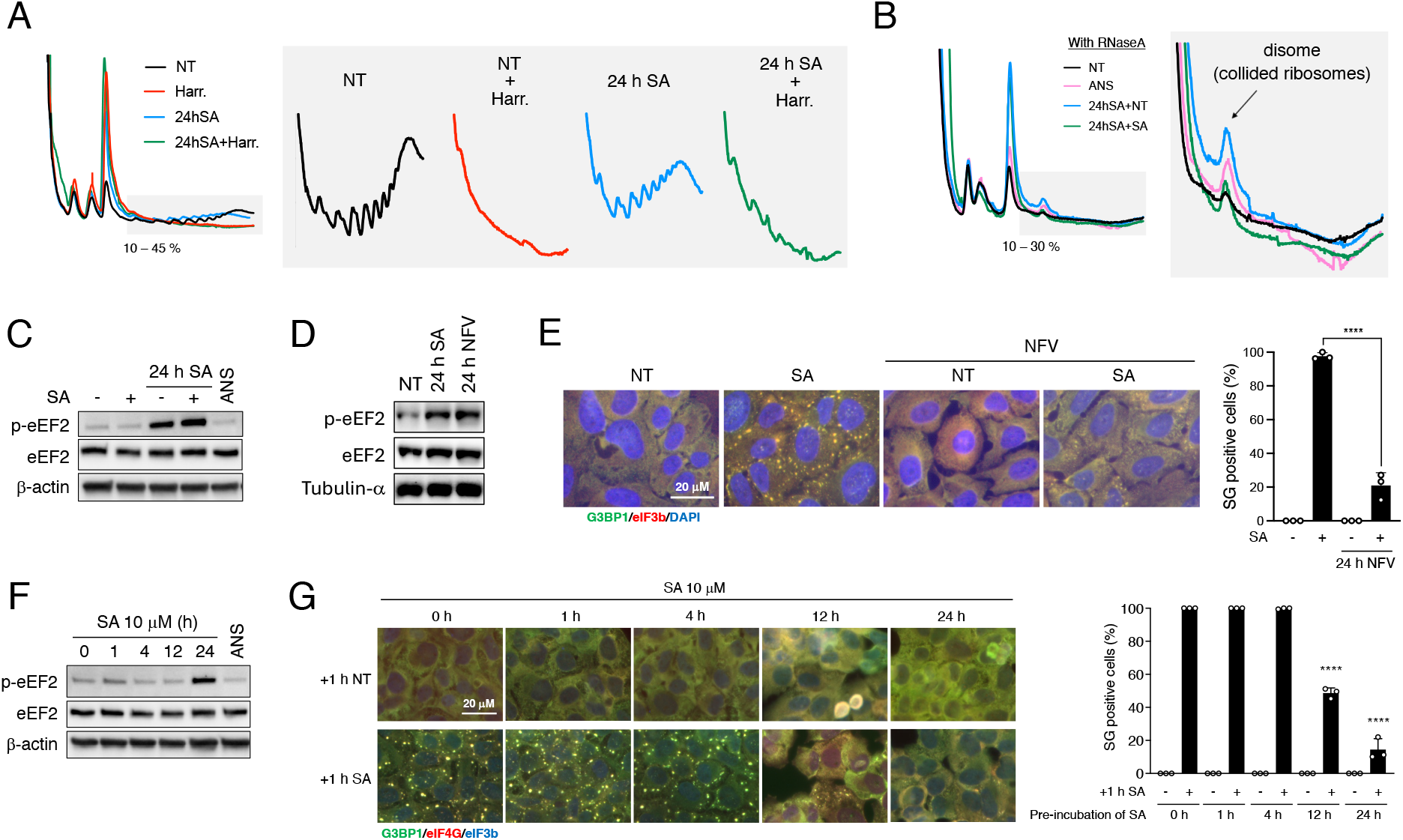
Chronic stress slows down translation at the elongation step. (A) Polysome profiles from U2OS cells. NT; black, 10 min of 2 μM Harringtonin (Harr.); red, 24 h 10 μM SA pre-incubation; blue, 10 min of 2 μM Harringtonin (Harr.) with 24 h 10 μM SA pre-incubation; green. (B) Polysome profiles from RNase A (0.5 mg/ml)-digested lysates of U2OS cells. NT; black, 1 mg/ml ANS; pink, 10 μM SA incubation for 24 h; blue, 100 μM SA treatment for 1 h with 10 μM SA pre-incubation for 24 h; green. (C) U2OS cells were subjected to treatment with 100 μM SA for 1 h after pre-incubation with 10 μM SA, or 1 mg/ml ANS for 15 min. Cell lysates were subjected to western blotting using antibodies for p-eEF2, total eEF2, and β-actin. (D) U2OS cells were subjected to treatment with 10 μM SA, or 12.5 μM NFV (Nelfinavir) for 24 h. Cell lysates were subjected to western blotting using antibodies for p-eEF2, total eEF2, and α-tubulin. (E) U2OS cells were subjected to treatment with 100 μM SA for 1 h after pre-incubation with 12.5 μM NFV for 24 h. Cells were examined for the presence of the core SG markers G3BP1 (green), eIF3b (red), and DAPI (blue). G3BP1 and eIF3b positive cells were quantified. Results are mean ± S.E.M. (n = 3). P values were assessed using a one-way ANOVA (*p*\*\*\*\* < 0.0001) (F) U2OS cells were subjected to treatment with 10 μM SA for 1, 4, 12, 24 h or 1 mg/ml ANS for 15 min. Cell lysates were subjected to western blotting using antibodies for p-eEF2, total eEF2, and β-actin. (G) U2OS cells were subjected to treatment with 100 μM SA for 1 h after pre-incubation with 10 μM SA for 0, 1, 4, 12, and 24 h. Cells were examined for the presence of the core SG markers G3BP1 (green), eIF4G (red), and eIF3b (blue). All positive cells were quantified. Results are mean ± S.E.M. (n = 3). P values were assessed using a one-way ANOVA (vs. 1 h Pre-incubation of SA +1 h SA; *p*\*\*\*\* < 0.0001)

While moderate ribosome collisions are efficiently rescued by the RQC response program, an excessive ribosome collision activates the stress-sensing MAP3 kinase ZAKα, which binds ribosomes and acts as one of the arms of RSR (24–27). At low doses, (1 mg/ml) ANS is known to induce ribosome collision, whereas higher doses freeze ribosome elongation without inducing collisions (24). We used polysome profiling followed by RNaseA treatment to quantify the ribosome fraction involved in stalling or collisions (manifest by disomes) (**Fig. 5B**). Chronic stress pre-conditioning (24 h SA + NT and 24 h SA + 1 h SA) produces polysome peaks similar to those observed in cells treated with 1 mg/ml ANS for 15 min. We confirmed that the observed peaks are reminiscent of disomes of collided ribosomes (as in control ANS-treated sample), which are resolved to monosomes following exposure to a higher dose of RNase (**Sup Fig. 6 A**).

To determine whether stalled ribosomes actually represent collided ribosomes, we used phos-tag SDS PAGE to quantify ZAKα phosphorylation in chronic SA treated cells (**Sup Fig. 6 B – C**). Our data suggest that 24 h low dose SA treatment only partially triggers ZAKα phosphorylation. In addition, we determined whether chronic SA treatment will promote ubiquitination of RPS10, a ribosomal protein that is specifically modified during excessive ribosome collisions such as promoted by ANS treatment (37, 38) (**Sup Fig. 6 D**). Our data suggest that stalled ribosomes mostly do not represent bona fide collided ribosomes, which direct are targets of RSR. Thus, chronic stress induces ribosome stalling that is different to excessive collisions observed during excessive ribotoxic damage.

We further analyzed the eukaryotic elongation factor 2 (eEF2) phosphorylation status. eEF2 regulates translation elongation by GTP-dependent translocation of peptidyl-tRNA, but phospho-eEF2 inhibits this step and translation, leading to ribosome stalling (22). Indeed, low dose SA pre-conditioning induced phosphorylation of eEF2 while ANS does not (**Fig. 5 C**). As Nelfinavir (NFV) is known to phosphorylate eEF2 (39), we incubated cells with NFV chronically for 24 h and checked the ability to induce SGs by additional SA incubation. We observed that NFV induced same level of phosphorylation of eEF2 as 24 h 10 μM SA, and significantly inhibited SG formation by additional acute stress (**Fig. 5 D – E**).

Next, we examined the profiles of phosphorylation of ZAKα and eEF2 in a time-dependent manner. While phosphorylated ZAKα was only partially observed at both 12 and 24 h (with phosphorylation stronger at 24 h than 12 h) (**Sup Fig. 6 D**), eEF2 was only significantly phosphorylated at 24 h (**Fig. 5 F**). These results indicate the phospho-eEF2-induced translocation failure may slowly accumulate in form of stalled ribosomes which may lead to severe ribosome collision at much lates stages, but it is not the trigger of collision itself. We also determined the ability of cells to form SGs (by an addition of 100 μM SA) at each time point. The percentage of SG-positive cells decreased in 12h and 24 h pre-incubated cells, coinciding with levels of ZAKα phosphorylation (**Fig. 5 G**).

These data suggest that chronic stress pre-conditioning slows down mRNA translation, promotes ribosome stalling, which consequently delay polysome disassembly and SG formation in response to additional acute stress (**Fig. 6**).

**Figure 6.**
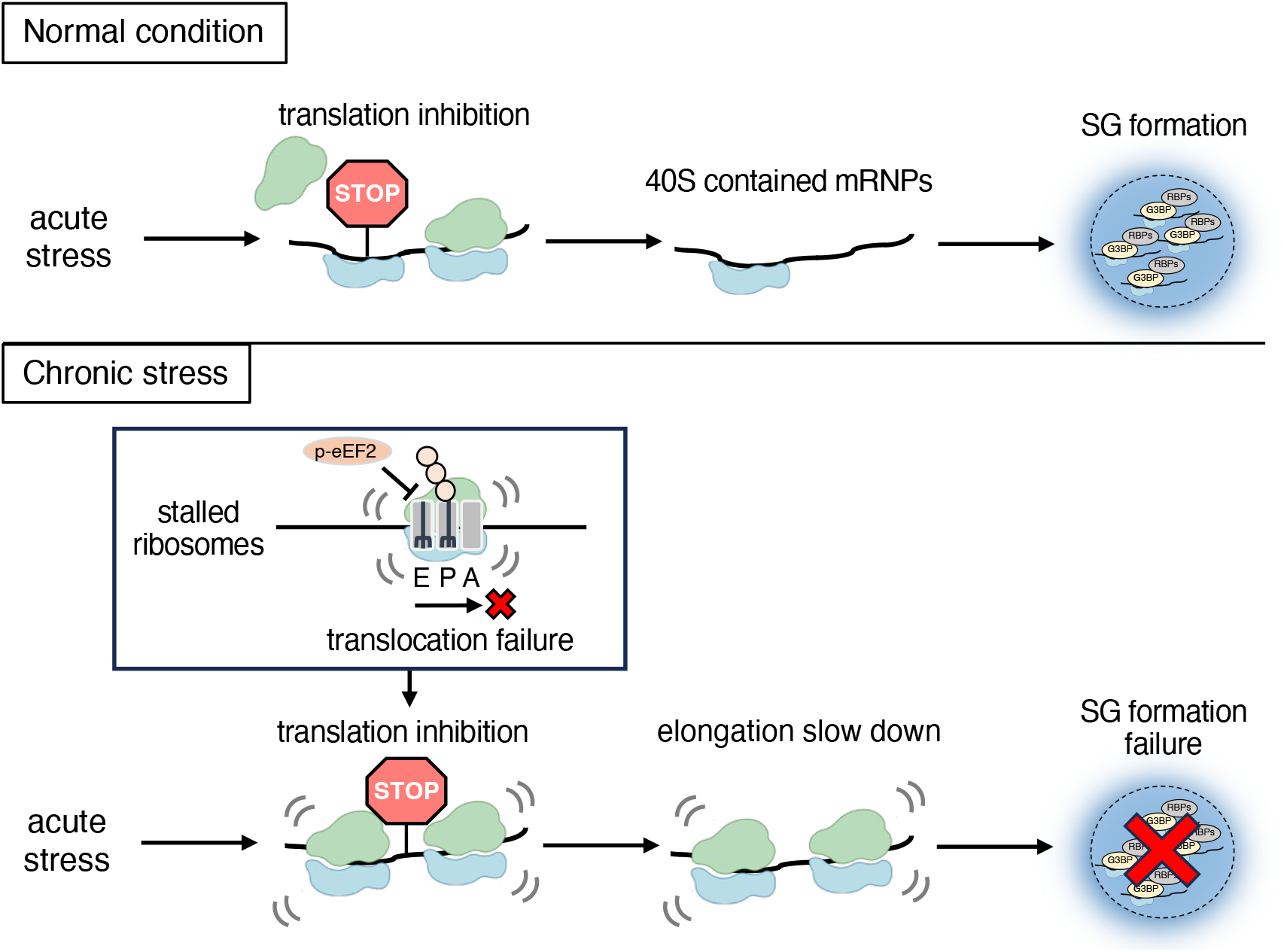
Schematic illustration of SGs formation failure by translation slowdown in chronic stress pre-incubated cells. Acute SA stress induces translation inhibition thereby splitting ribosomes, which inducts SG formation. Chronic stress incubation makes ribosomes stalled with translocation failure, which induces translation slowdown by additional acute stress despite translation inhibition, and SG formation failure as a result.

## Discussion

SG assembly and disassembly in response to acute stress has been extensively studied (40). SG dynamics following chronic stress has received less attention, despite SGs playing an important role in the pathophysiology of neurodegenerative disease, cancer, inflammatory disease, and other chronic stress-related conditions (8, 14, 41, 42). While we showed that any regime of SA-induced chronic stress itself did not induce SG formation (**Fig. 1 A**), it has been reported that chronic incubation (30 h) of low-dose (15 μM) SA induces SG formation in iPSC-motoneurons (43) suggesting cell type-specific differences in response to stresses. Interestingly, aging is considered as a condition of complex chronic stress that induces cellular senescence, a stable cell cycle arrest linked to various diseases (44–46). Since it is known that aging/cellular senescence promotes ribosome collision and translational repression (26, 47–49), our data suggest that aged cells may have a reduced propensity to form SGs, a phenotype that may be intrinsically linked to chronic stress-induced changes in gene expression.

Our studies indicate that most of chronic stress conditions inhibit both initiation (judged by eIF2α phosphorylation) and elongation (eEF2 phosphorylation) phases of protein synthesis while acute stress tends to target the initiation step. We speculate here that slow down of translation by eEF2 phosphorylation under conditions of chronic stress aims on the maintenance of cellular metabolism mostly by energy conservation, and by making slight adjustments in gene expression. This is in contrast to extensive stress (e.g., causing ribosome collision and activation of RSR) or transient acute stress (causing abrupt global translation arrest coupled with sequestration of mRNPs into SGs), which aim on cell survival and make drastic changes in gene expression.

Mechanistically, G3BP1/2 (G3BP) proteins are SG nucleators that are required for SG formation under most of stresses, except osmotic. As G3BP needs to bind 40S ribosomal subunits to nucleate SGs, 80S ribosome dissociation and polysome disassembly both contribute to SG formation (29). Although our results suggest that ribosome stalling and inhibition of translation elongation inhibits SG disassembly in chronic stress pre-conditioned cells, distinct polysome profiles are observed in cells exposed to different types of stress (**Fig. 2 B – E**, blue and green), whereas the percentage of SG-positive cells is almost the same in each case (**Fig. 1 C – F**). One potential explanation is that the presence of a single 80S ribosome on mRNA is sufficient to prevent its recruitment into SGs, as suggested by (50). Puromycin is known to promote SG formation via polysome disassembly (**Sup Fig. 5)** (15), but chronic stress pre-conditioned cells show only partial disassembly of polysomes in response to puromycin (**Fig. 4 B**). It should however be noted that high doses of SA are able to promote SG formation suggesting that SG components such as pool of mRNAs (**Sup Fig. 4)** and SGs nucleators (**Fig. 2 A**) are present in sufficient amounts to trigger SG formation per se. In agreement with this notion, we could not detect any significant differences in poly(A) mRNA levels or proteins playing roles in SG dynamics.

Our analysis of the effects of chronic stress on polysome formation suggests that stalled ribosomes contribute to changes in translation and SG assembly. We also showed that chronic stress-induced eEF2 phosphorylation may promote ribosome stalling via translocation failure, but does not trigger extensive collisions, at least at the earlier time points of stress exposure (**Sup Fig. 6B – C, Fig. 5 C**). The exact contribution of chronic stress-induced ribosome collisions in the observed slowdown of translation is still unclear but there are several possibilities: an insufficient supply of aminoacyl-tRNAs, translation of damaged mRNAs, damaged ribosomes, etc. (19–21). Slowing down translation also can be beneficial for correct protein folding and oppose proteostasis, a hallmark of chronically stressed cells. Ribosome stalling and eEF2 phosphorylation is not increased abruptly but increased in a time-dependent manner (**Fig. 5 C, Sup Fig 6**), which provide RQC enough time to adjust and be efficient under chronic stress. Loss of function experiments of RQC proteins, which are observed under excessive stress, suggest that RQC failure is a trigger of cell death, which can contribute to some diseases (51, 52). Our data also indicate that the inability to promote SGs may also contribute to cell death.

In conclusion, as summarized in **Fig. 6**, we propose that chronic stress-induced eEF2 phosphorylation slows down mRNA translation, causing ribosome stalling, which in turn inhibits SG formation. Inefficient disassociation of stalled 80S ribosomal subunits may directly contribute to this phenomenon. While canonical SGs are proposed to have a cytoprotective role (5), chronic stress-induced failure of SG formation together with other processes such as RQC mechanisms may activate cell death. Ribosome stalling and collisions are known to induce proteostasis that ultimately contributes to human diseases via cell death pathways (25, 26, 28). Our data have uncovered a new functional connection between ribosome stalling and SG formation under chronic stress and provide a foundation for understanding and exploring acute stress in the context of chronic stress models such as aging.

## Materials and Methods

### Cell culture and drug treatment

Human osteosarcoma U2OS cells were maintained at 37 °C in a 5.0% CO2 in DMEM (Corning) containing 20 mM HEPES (Gibco), 10% FBS (Sigma), 100 U/ml penicillin, and 100 μg/ml streptomycin. For any experiments, cells were grown to ∼70% confluency and then treated as indicated in figure legends: sodium arsenite (SA, Sigma), rocaglamide A (RocA, MedChemExpress), menadion (Mena, Sigma), doxorubicin (Drb, AdooQ Bioscience), thapsigargin (Tg, Invitrogen), tunicamycin (Tun, Sigma), brefeldin A (BA, Sigma), carbonyl cyanide 3-chlorophenylhydrazone (CCCP, Sigma), 2-deoxy-D-glucose (2DG, Sigma), nelfinavir (NFV, MedChemExpress), arsenic trioxide (ATO, Sigma). Puromycin (puro, Sigma) treatment was performed 30 min before collecting the coverslips or as described in the text. Anisomycin (ANS, MedChemExpress) was performed 15 min before harvest or 15 min before the incubation with SA as indicated.

### Western blotting

Following drug treatment, cells were washed with phosphate-buffered saline (PBS). Protein samples were heated to 95°C for 10 min in Laemmli sample buffer in the presence of 100 mM dithiothreitol (DTT) and subjected to SDS–PAGE. Samples were loaded on a 4–20% Tris-Glycine gel (BioRad) and transferred to nitrocellulose membrane. Membranes were blocked with Tris-buffered saline with 0.1% Tween-20 (TBS-T) with 5% milk for 1 h at room temperature. Antibodies were diluted in 5% normal horse serum in PBS. Primary antibodies were incubated overnight at 4°C and secondary antibodies for 1 h at room temperature. Antibody information is listed in Table S1. Antibody detection was performed using SuperSignal West Pico Chemiluminescent Substrate (Thermo Scientific).

### RiboPuromycylation assay

Ribopuromycylation assay was described in (53). In brief, cells were unstressed or stressed as indicated. 5 min before harvest, puromycin and emetine were added to a final concentration of 9 and 91 μM, respectively, and the incubation continued for 5 min. Cells were then lysed and subjected to Western blotting using an anti-puromycin antibody.

### M^7^GTP pull-down assay

Cells were lysed by lysis buffer (Tris–HCl pH7.4, 100 mM NaCl, 0.5% NP-40, protease inhibitor (Thermo Scientific)), and centrifuged for 15 min at 12,000×g at 4°C. The supernatant was mixed with Immobilized g-aminophenyle-m7gtp (Jena Bioscience) for 2 h at 4°C with rotation. The beads were washed extensively with the lysis buffer and cap-bound materials were eluted by boiling in Laemmli sample buffer supplemented with 100 mM DTT.

### mRNA pull-down, cDNA synthesis, and qPCR

Total RNA was extracted by using Trizol (Invitrogen). mRNA was extracted by polyA+ purification with Dynabeads™ mRNA DIRECT™ Purification Kit (Ambion). All RNAs were measured by NanoDrop™ One (thermo fisher). Both 30 ng of total RNA and pulled-down mRNA were reverse transcribed with the Superscript IV first-strand synthesis kit for RT-qPCR (ThermoFisher). The qRT-PCR was performed by using iQ™ SYBR® Green Supermix (Bio-Rad), cDNA template, and gene-specific primer sets designed using the IDT primer design tool. No reverse transcriptase and no template controls were performed in parallel to check for DNA contamination and primer-dimer. The primer sets used for the study are given in Table S2. Threshold cycle (CT) values in qRT-PCR experiments were averaged across three biological replicates.

### Immunofluorescence

Cells were plated into a 24-well plate seed with coverslips. The following day, cells were treated as indicated in figure legends. Then the cells were fixed with 4% paraformaldehyde for 15 min, permeabilized with −20 °C methanol for 5 min, and blocked for 1 h with 5% normal horse serum (NHS; ThermoFisher) diluted in PBS. Primary antibodies shown in Table S1 were diluted in blocking solution and incubated for 1 h at room temperature or overnight at 4 °C. Next, cells were washed three times and then incubated with secondary antibodies (Jackson Laboratories) and Hoechst 33258 (Sigma-Aldrich) for 1 h at room temperature and washed. Coverslips were mounted on glass slides with Vinol and imaged.

### Fluorescence in situ hybridization (FISH)

For in situ hybridization, cells were fixed with 4% paraformaldehyde for 15 min and then permeabilized with −20°C methanol for 5 min. Cells were incubated overnight in 70% ethanol at 4°C. The following day, cells were washed twice with 2× saline-sodium citrate (SSC), blocked in hybridization buffer (Sigma) for 30 min, then hybridization was performed using a biotinylated oligo(dT40) probe (2 ng/μl) diluted in hybridization buffer at 37°C. After extensive washes with 2× SSC at 37°C the probe was revealed using Cy-conjugated streptavidin (Jackson Immunoresearch Laboratories), followed by immunostaining as described above.

### Microscopy

Wide-field fluorescence microscopy was performed using an Eclipse E800 microscope (Nikon, Minato, Tokyo, Japan) equipped with epifluorescence optics and a digital camera (Spot Pursuit USB). Image acquisition was done with a 40× objective (PlanApo; Nikon, Minato, Tokyo, Japan). Images were merged using Adobe Photoshop.

### Polysome profiling

Cells were treated with 100 mg/ml cycloheximide (Sigma) for 10 min, washed with PBS, and harvested into lysis buffer (10 mM Tris [pH 7.4], 150 mM NaCl, 5 mM MgCl2, 1 mM DTT, 100 μg/ml cycloheximide, 1% Triton-X100) supplemented with RNasin Plus inhibitor (Promega) and HALT phosphatase and protease inhibitors (Thermo Scientific). Lysates were rotated at 4 °C for 10 min, and centrifuged for 5 min at 12,000×g. Supernatants were loaded onto 10 – 45 % sucrose gradients made in gradient buffer and centrifuged in a Beckman SW 40 Ti rotor for 3.5 h, 29,000xg at 4 °C. Samples were eluted and its OD 254 was measured by a Biocomp gradient station attached to a syringe pump. With RNase experiments, supernatants of lysates were incubated with RNase A (Invitrogen) at 37 °C for 15 min, then loaded onto 10 – 30% sucrose gradients.

## Acknowledgments

We thank members of Ivanov and Anderson labs for the valuable discussion. The figures are created with BioRender.com. This work was supported by funds from National Institutes of Health grant R35 GM126901 (P.J.A.), National Institutes of Health grant R01 GM126150 and R01 GM146997 (P.I.), Japan Society for the Promotion of Science (JSPS) Grants-in-Aid for Scientific Research (PS KAKENHI) 22KJ2354 (Y.A.), 20K21761 and 21H03359 (M.M.), 19H04053 and 23H03329 (Y.T.). Y.A. was supported by the JSPS Overseas Challenge Program for Young Researchers.

