## Supplementary material for "Chronic stress antagonizes formation of Stress Granules": Supp Information

### Supplementary Table 1. List of antibodies

| Name | Species | Source | Identifer | Dillusion | Experiment |
|---|---|---|---|---|---|
| **α-tubulin** | Rabbit | Proteintech | 11224-1-AP | 1/3000. | WB |
| **β-actin** | Mouse | Proteintech | 66009-1-lg | 1/5000. | WB |
| **Caprin1** | Rabbit | Proteintech | 15112-1-AP | 1/1000. | WB |
| **eEF2** | Rabbit | Cell Signaling Technology | #2332 | 1/1000. | WB |
| **eIF3b** | Goat | Santa Cruz Biotechnology Inc. | sc-16377 | 1/250. | IF |
| **eIF4A** | Mouse | Santa Cruz Biotechnology Inc. | sc-377315 | 1/500. | WB |
| **eIF4E** | Rabbit | Cell Signaling Technology | #9742 | 1/1000. | WB |
| **eIF4G** | Rabbit | Santa Cruz Biotechnology Inc. | sc-11373 | 1/250 for IF, 1/1000 for WB. | IF, WB |
| **G3BP1** | Mouse | Santa Cruz Biotechnology Inc. | sc-81940 | 1/250 for IF, 1/1000 for WB. | IF, WB |
| **G3BP2** | Rabbit | Bethyl Laboratories Inc. | A302-040A | 1/1000. | WB |
| **Phospho-eEF2 (Thr56)** | Rabbit | Cell Signaling Technology | #2331 | 1/1000. | WB |
| **Phospho-eIF2α (Ser51)** | Rabbit | Cell Signaling Technology | #9721 | 1/2000. | WB |
| **puromycin** | Mouse | Sigma-Aldrich | MABE343 | 1/1000. | WB |
| **total eIF2α** | Mouse | Cell Signaling Technology | L57A5 | 1/2000. | WB |
| **USP10** | Rabbit | Bethyl Laboratories Inc. | A300-900A | 1/1000. | WB |
| **ZAKa** | Rabbit | Bethyl Laboratories Inc. | A301–933A | 1/1000. | WB |
| **RPS10** | Rabbit | Abcam | ab151550 | 1/500 | WB |

### Supplementary Table 2. List of primers

| species | gene name | direction | sequence (5'-) |
|---|---|---|---|
| **Human** | GAPDH | F | ACATCGCTCAGACACCATG |
| | | R | TGTAGTTGAGGTCAATGAAGGG |
| | AHNAK | F | TTCAGTCTCCACCCCAAATG |
| | | R | GAACAATGCTCCAAAGAACGG |
| | DYNC1H1 | F | ACCATCGTCAACTTCTCTGC |
| | | R | CAATTGCCCATTTCACCATAGG |
| | NORAD<br>(non cording RNA: LINC00657) | F | AAGCTGCTCTCAACTCCACC |
| | | R | GGACGTATCGCTTCCAGAGG |
| | BACTIN | F | GGAAATCGTGCGTGACATTAAG |
| | | R | TAGTCCGCCTAGAAGCATTTG |
| | 18S rRNA | F | ACGTCTGCCCTATCAACTTTC |
| | | R | CCGCGGTCCTATTCCATTATT |
| | 28S rRNA | F | CGGGATAAGGATTGGCTCTAAG |
| | | R | CTGTGGTTTCGCTGGATAGT |

### Supplementary Figure 1.

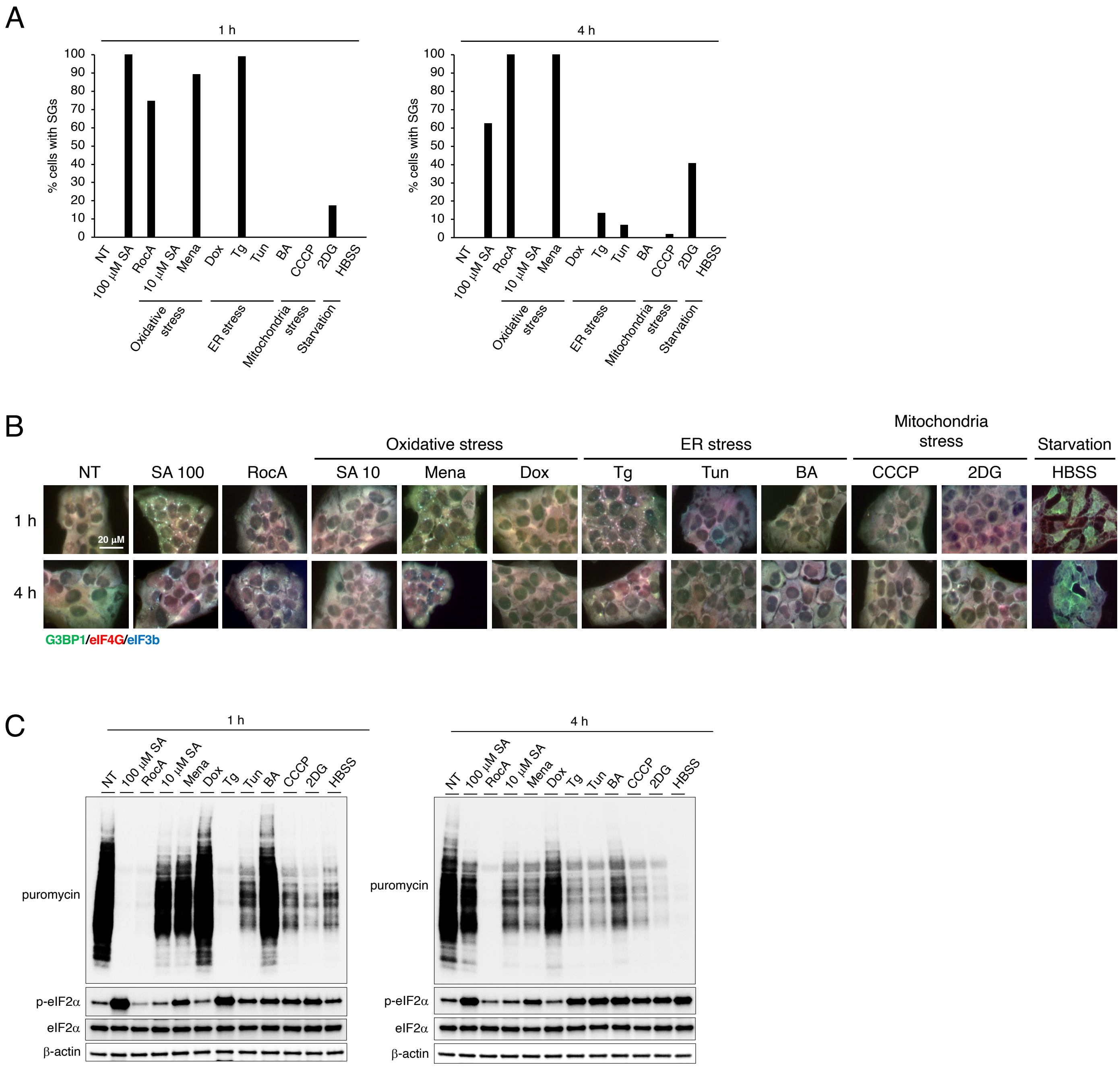

### Supplementary Figure 2.

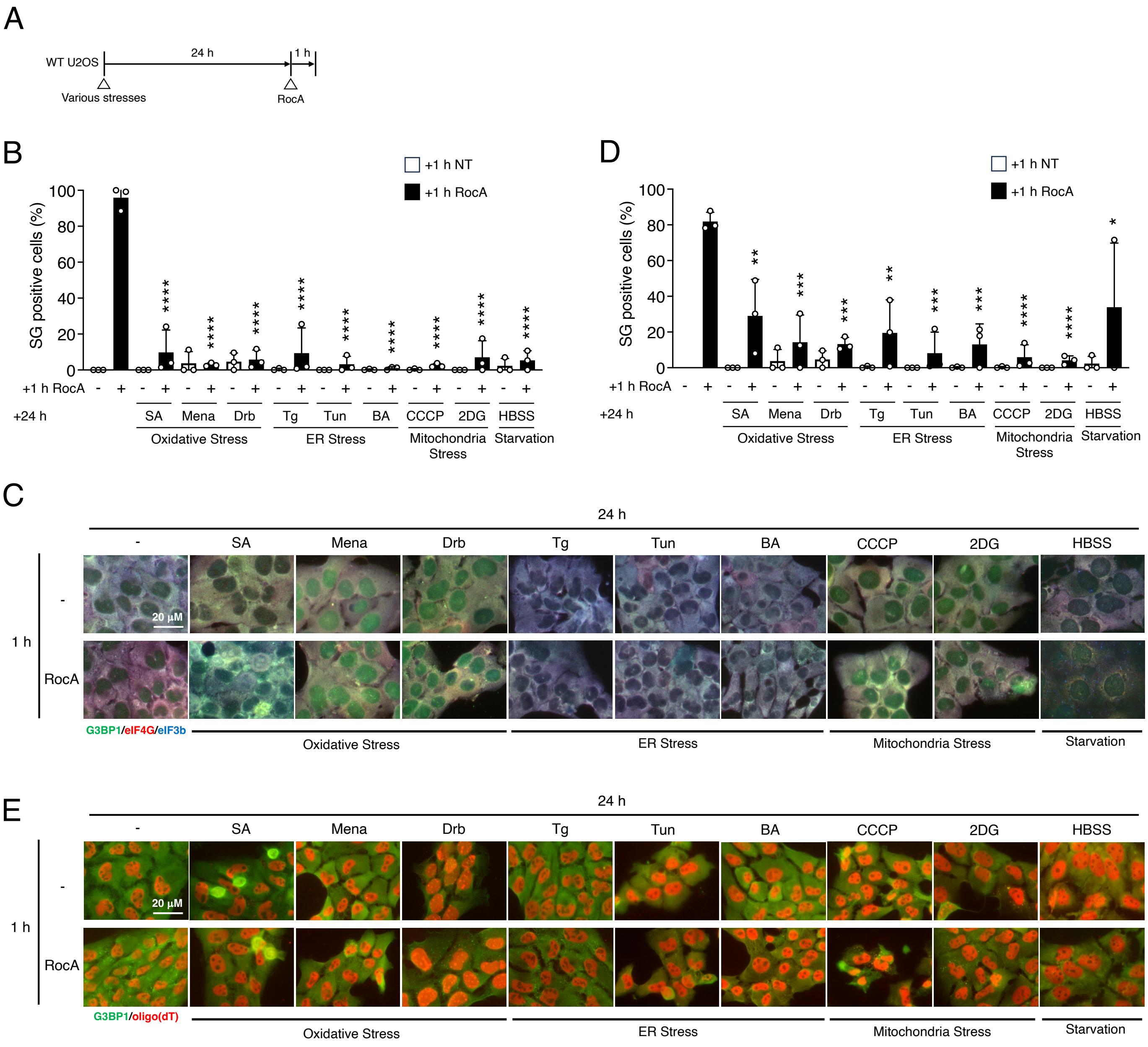

### Supplementary Figure 3.

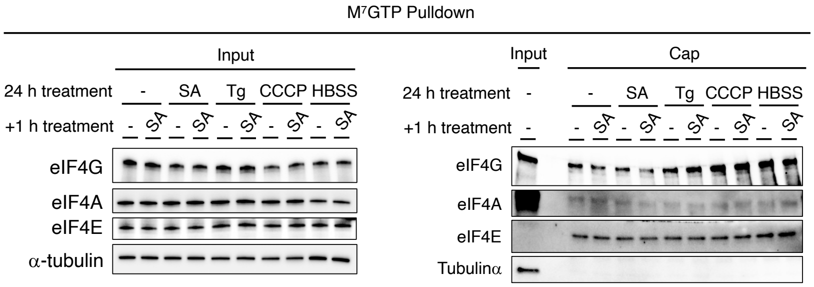

### Supplementary Figure 4.

A

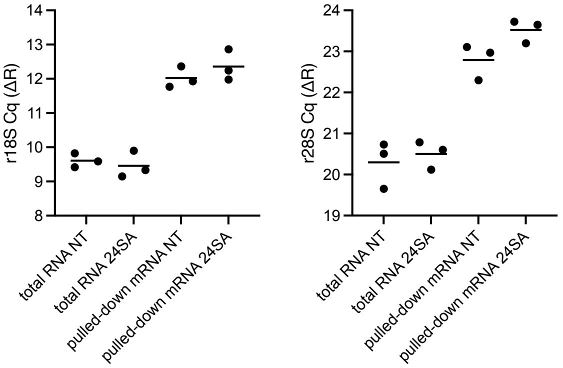


B

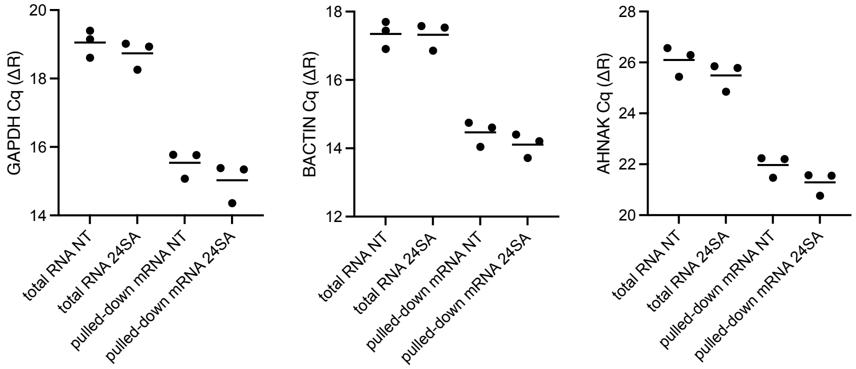

### Supplementary Figure 5.

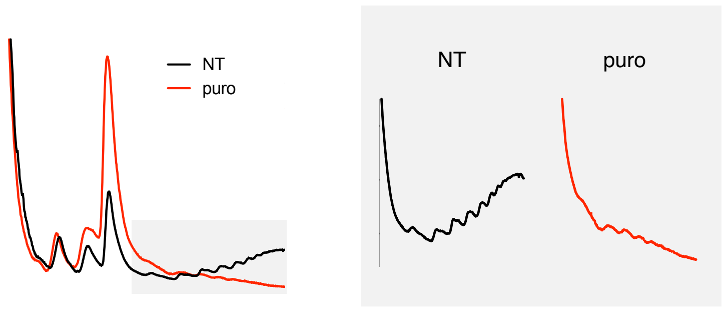

#### Supplementary Figure 6.

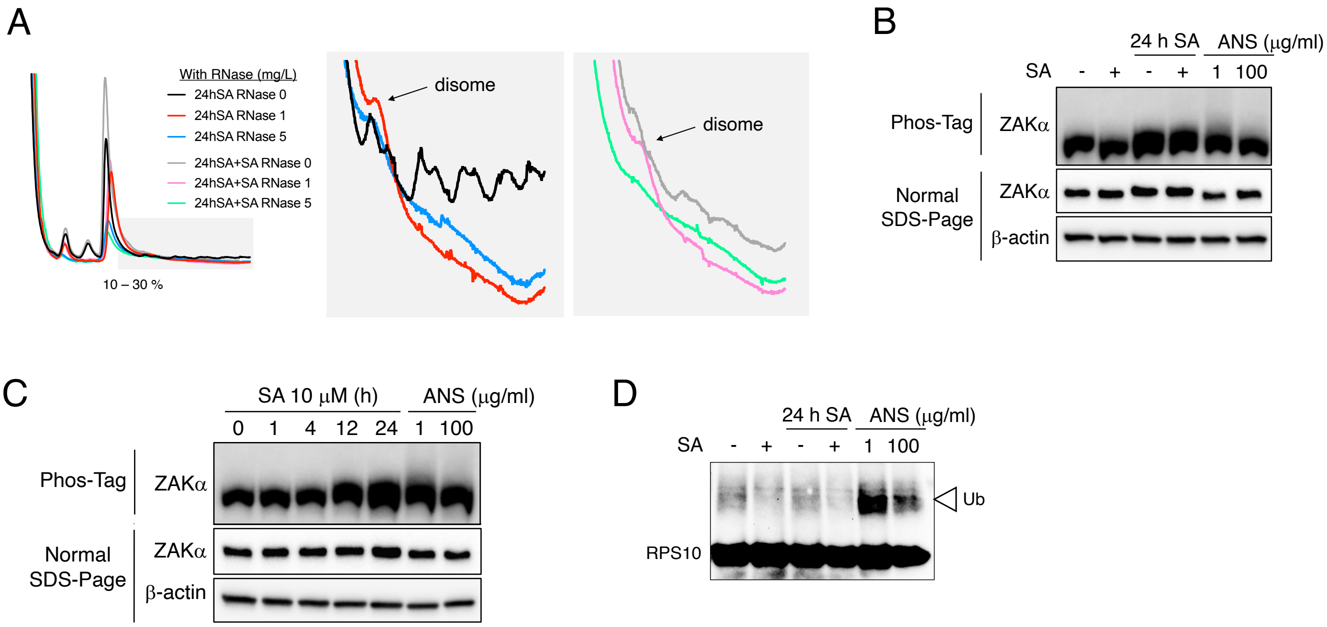

**Supplementary information**

**Supplementary Figure 1. Sg response to several acute stress inducers**

U2OS cells were subjected to treatment with 100 $\mu$M SA, 2 $\mu$M RocA (rocaglamide A), 10 $\mu$M SA, 30 $\mu$M Mena (menadione), 2 $\mu$M Drb (doxorubicin), 1 $\mu$M Tg (thapsigargin), 25 $\mu$g/ml Tun (tunicamycin), 25 $\mu$g/ml BA (brefeldin A), 60 $\mu$M CCCP (carbonyl cyanide m-chlorophenyl hydrazone), 60 mM 2DG (2-deoxy-D-glucose), or HBSS for 1 or 4 h. Unstressed cells (NT) were used as a control. (A) Cells were examined for the presence of the core SG markers G3BP1, eIF4G, and eIF3B. (B) Representative images of U2OS cells stained with G3BP1 (green), eIF4G (red), and eIF3B (blue) after the cells had been subjected to specific stresses. (C) Cells were pulsed with puromycin and emetine for 5 min and lysed. Cell lysates were subjected to western blotting using antibodies for Puromycin, p-eIF2α, total eIF2α, and β-actin.

**Supplementary Figure 2. Acute RocA treatment in chronic stress pre-incubated cells**

(A) Schematic illustration of the experimental timeline (B–E) U2OS cells were subjected to treatment with 2 $\mu$M RocA for 1 h after preincubation with 10 $\mu$M SA, 30 $\mu$M Mena (menadione), 2 $\mu$M Drb (doxorubicin), 1 $\mu$M Tg (thapsigargin), 25 $\mu$g/ml Tun (tunicamycin), 25 $\mu$g/ml BA (brefeldin A), 60 $\mu$M CCCP (carbonyl cyanide m-chlorophenyl hydrazone), 60 mM 2DG (2-deoxy-D-glucose), or HBSS for 24 h. Unstressed cells (NT) were used as a control. (B) Cells were examined for the presence of the core SG markers G3BP1, eIF4G, and eIF3B. (C) Representative images of U2OS cells stained with G3BP1 (green), eIF4G (red), and eIF3B (blue) after the cells had been subjected to the specific stresses. (D) Cells were examined for the presence of the core SG marker G3BP1 and poly (A) mRNAs [FISH using oligo(dT) probe]. (E) Representative images of U2OS cells stained with G3BP1 (green) and oligo(dT) (red) after the cells had been subjected to specific stresses. (B and D) P values were assessed using a one-way ANOVA (vs. 1 h RocA; $p^{****} < 0.0001$, $p^{***} < 0.001$, $p^{**} < 0.01$, $p^{*} < 0.05$) Results are mean ± S.E.M. (n = 3). NT samples are represented in Fig. 1.

**Supplementary Figure 3. Analysis of eIF4F complex integrity by m$^7$GTP pulldown under chronic stress pre-incubation**

U2OS cells were subjected to treatment with 100 $\mu$M SA for 1 h after pre-incubation with 10 $\mu$M SA, 1 $\mu$M Tg, 60 $\mu$M CCCP, or HBSS for 24 h. Unstressed cells (NT) were used as a control. Cells were lysed and subjected to m$^7$GTP-sepharose pulldown to isolate cap-associated proteins. Both the input and the precipitate were processed for western blotting and probed for the presence of eIF4E, eIF4G, eIF4A and α-tubulin.

**Supplementary Figure 4. qPCR detection of SS-associated and previously validated target mRNAs**

U2OS cells were incubated with 10 μM of SA for 24 h (24SA), or NT and total RNA was extracted. mRNA was pulled down from total RNA, 30 ng of both RNAs were reverse transcribed and qRT-PCR was performed. (A) Threshold cycle (CT) values in qRT-PCR experiments of ribosomal RNA 18 S and 28 S. (B) CT values in qRT-PCR experiments of GAPDH, BACTIN, and AHNAK. CT values were averaged across three biological replicates.

**Supplementary Figure 5. Polysome profiling of puromycin incubation (control experiment)**

Polysome profiles from U2OS cells. NT; black, 20 μg/ml of puromycin (puro) for 0.5 h; red.

**Supplementary Figure 6. Chronic stress may induce partial ribosome collisions**

(A) Polysome profiles from RNase A digested lysates of U2OS cells. Cells were incubated with or without 100 μM SA for 1 h, with 10 μM SA pre-incubation for 24 h. 24 h SA with RNase 0 mg/ml; black, 24 h SA with RNase 1 mg/ml; red, 24 h SA with RNase 5 mg/ml; blue, 24 h SA + 1 h SA with RNase 0 mg/ml; gray, 24 h SA + 1 h SA with RNase 1 mg/ml; pink, 24 h SA + 1 h SA with RNase 5 mg/ml; light green. (B) Cell lysates were subjected to western blotting by Phos-tag gels for ZAKα phosphorylation in U2OS cells treated with 100 μM SA for 1 h after pre-incubation with 10 μM SA or 15 min ANS at 1, 100 mg/ml. (C) Cell lysates were subjected to western blotting by Phos-tag gels for ZAKα phosphorylation in U2OS cells treated with 10 μM SA for 1, 4, 12, 24 h or 15 min ANS at 1, 100 mg/ml. (D) Cell lysates were subjected to western blotting in U2OS cells treated with 100 μM SA for 1 h after pre-incubation with 10 μM SA or 15 min ANS at 1, 100 mg/ml. Cell lysates were subjected to western blotting using antibodies for RPS10. Ub: Ubiquitinated-RPS10 band.
